# A national analysis of trends in COVID-19 infection and clinical management in Veterans Health Administration medical facilities

**DOI:** 10.1101/2021.01.18.427092

**Authors:** Maya Aboumrad, Brian Shiner, Natalie Riblet, Hugh Huizenga, Nabin Neupane, Yinong Young-Xu

**Author notes:** **Corresponding Author:** Maya Aboumrad, Veterans Affairs Medical Center, 215 North Main Street, White River Junction, VT 05009.

## Abstract

**OBJECTIVE:** We explored longitudinal trends in sociodemographic characteristics, reported symptoms, laboratory findings, pharmacological and non-pharmacological treatment, comorbidities, and 30-day in-hospital mortality among hospitalized patients with coronavirus disease 2019 (COVID-19).

**METHODS:** This retrospective cohort study included 43,267 patients diagnosed with COVID-19 in the Veterans Health Administration between 03/01/20 and 08/31/20 and followed until 09/30/20. We focused our analysis on patients that were subsequently hospitalized, and categorized them into groups based on the month of hospitalization. We summarized our findings through descriptive statistics. We used a nonparametric rank-sum test for trend to examine any differences in the distribution of our study variables across the six months.

**RESULTS:** During our study period, 8,240 patients were hospitalized, and 1,081 (13.1%) died within 30 days of admission. Hospitalizations increased over time, but the proportion of patients that died consistently declined from March (N=221/890, 24.8%) to August (N=111/1,396, 8.0%). Patients hospitalized in March compared to August were younger on average, mostly black, and symptomatic. They also had a higher frequency of baseline comorbidities, including hypertension and diabetes, and were more likely to present with abnormal laboratory findings including low lymphocyte counts and elevated creatinine. Lastly, receipt of mechanical ventilation and Hydroxychloroquine declined from March to August, while treatment with Dexamethasone and Remdesivir increased.

**CONCLUSION:** We found evidence of declining COVID-19 severity and fatality over time within a national health care system.

## INTRODUCTION

Coronavirus disease 2019 (COVID-19) is an infectious respiratory illness caused by the severe acute respiratory syndrome coronavirus 2 (SARS-CoV-2).^1,2^ As of November 19, 2020, the United States (U.S.) has the highest number of reported COVID-19 cases (11.5 million) and deaths (249,670) globally.^2^ There is a growing body of literature utilizing electronic medical record data to provide “real-world” information on risk factors for COVID-19 infection, severity, and fatality including certain sociodemographic characteristics, symptoms, laboratory findings, underlying comorbidities, and treatments.^1,3–17^ However, the results of these studies are largely conflicting, especially among those conducted during the initial onset of the pandemic compared to more recent literature, as well as among those that were regional or local compared to national in scope.^1,3–18^

Conflicting findings from “real-world” studies may be at least partially due to lack of consideration for the temporal and regional distribution of COVID-19 burden across the U.S., secular trends in clinical practice, or potential systemic differences in the characteristics of patients requiring hospitalization or treatment.^9,13,15,16,18,19^ Moreover, the aggregation of this information at one time point and short-term assessment (e.g., one month) may not accurately account for the influence of rapidly deployed and changing mitigation efforts including government restrictions and policies.^9,19–23^ Therefore, longitudinal assessment of changes over multiple time points would provide greater clarity on the epidemiology and clinical management of COVID-19 and allow for indirect assessment of mitigation efforts.

To address this knowledge gap, we explored potential trends and patterns in infection rates among all U.S. Veterans Health Administration (VHA) users diagnosed with COVID-19, and clinical management for those that were subsequently hospitalized at VHA medical facilities during the initial six months of the pandemic. Specifically, our objectives were to describe any changes from March 1, 2020 to August 30, 2020 in 1) sociodemographic characteristics, 2) reported symptoms, 3) laboratory findings, 4) pharmacological and non-pharmacological treatment, 5) 30-day in-hospital mortality, and 6) comorbidities among both hospitalized and in-hospital deceased patients. As the nation’s largest integrated health care system, the VHA was required to respond to COVID-19 in all geographic regions.^21,22^ Thus, the VHA offers a unique opportunity to better understand the case burden and management of COVID-19 due to its large size, diverse operating environment, nationally overseen policies and practices, and relatively homogenous patient population.^20–22^

## METHODS

This study received institutional review board approval from the Veteran’s Institutional Review Board of Northern New England at the White River Junction Veterans Affairs Medical Center.

### Data Source & Study Population

The VHA is comprised of over 170 medical centers and 1,250 community-based outpatient clinics.^24^ It provides comprehensive medical care to more than nine million veterans, including primary and specialty care.^24^ Additionally, VHA users have access to extensive inpatient care and treatment services including medical, surgical, mental health, dialysis, acute and long-term care.^24^ The VHA has an electronic medical record system with a centralized Corporate Data Warehouse (CDW), which contains longitudinal information on receipt of all services provided by VHA facilities including outpatient and inpatient visits, pharmacological and non-pharmacological treatments, and laboratory results, as well as patients’ sociodemographic and clinical characteristics. We obtained vital status from the VHA Vital Status File.

Our final cohort consisted of all VHA users (age ≥ 18 years) diagnosed with COVID-19 between March 1, 2020 and August 31, 2020 in accordance with Centers for Disease Control and Prevention standards, human confirmed case review, and/or VHA 10N memos.^20,25,26^

### Outcomes

Hospitalization: We limited our reporting to a subset of patients that were subsequently hospitalized for COVID-19 within the VHA health care system due to lack of complete clinical and outcome data for patients that received care at a non-VHA facility. Patients were indexed at the time of hospitalization for COVID-19 and categorized into six groups based on the month of hospitalization. We calculated hospital length of stay (LOS) as the number of days from admission to discharge or in-hospital death.

Mortality: Our primary endpoint of interest was 30-day in-hospital mortality. We examined mortality within 30 days of hospitalization to serve as a proxy for case fatality rate.^5^ We extended patient follow-up to September 30, 2020 to allow for sufficient assessment time of our study measures. Patients remained in the cohort until date of discharge or death.

### Study Variables

Patient Characteristics: We examined sociodemographic characteristics at the time of hospitalization for COVID-19 including age, sex, race/ethnicity, urbanicity of residence, and VHA priority rating (1-8). VHA priority group served as a proxy for socioeconomic status (SES) as it is partially based on income and the capacity for gainful employment. It is also connected to service-related disability, reflecting both health and VHA coverage.^27^ Priority group ratings range from 1-8, with group 1 representing the highest priority (i.e., lowest SES and complete healthcare coverage).

Symptoms: We examined COVID-related symptoms within 30 days preceding hospitalization for COVID-19 including abdominal pain, chills, common cold, cough, diarrhea, dyspnea, fatigue, fever, headache, myalgia, nausea, and sore throat.^6,14,28^

Laboratory Values: We examined laboratory values within seven days following hospitalization for COVID-19 including white blood cell count (WBC), absolute lymphocyte count (LC), platelet count (PC), creatinine, blood urea nitrogen (BUN), total bilirubin, aspartate aminotransferase (AST), alanine aminotransferase (ALT), ferritin, lactate, troponin I, brain-type natriuretic peptide (BNP), procalcitonin, c-reactive protein (CRP).^6,14,28^

Treatments: We examined receipt of any relevant/concomitant pharmacological and non-pharmacological treatment(s) during the course of patients’ hospitalization.

Pharmacological treatment included angiotensin-converting enzyme (ACE) inhibitors, antibiotics (e.g., Azithromycin), anticoagulants, antivirals (e.g., Hydroxychloroquine, Remdesivir), Azithromycin combined with Hydroxychloroquine, beta-blockers, bronchodilators, corticosteroids (e.g., Dexamethasone), immune-based therapy (e.g., Tocilizumab), non-steroidal anti-inflammatory drugs (NSAIDs), statins, and/or vasopressors.^29,30^ Non-pharmacological treatment included mechanical ventilation, dialysis, and supplemental oxygen.

Comorbidities: We examined the presence of clinical comorbidities within 12 months preceding hospitalization for COVID-19 according to the Charlson Comorbidity Index (CCI). The CCI score is a validated, weighted measure that predicts one-year mortality, with higher scores indicating greater illness burden.^31,32^ Additionally, we reported the top 10 comorbidities for both hospitalized and in-hospital deceased patients.

### Statistical Analysis

For all VHA users with laboratory-confirmed COVID-19, we reported the frequency of cases by month of diagnosis to examine trends in overall infection rates. For patients that were subsequently hospitalized for COVID-19 within the VHA, we reported our findings by month of hospitalization to examine trends in clinical management during the course of the pandemic.

We summarized our results through descriptive statistics. We reported proportion for categorical variables and mean (M) with standard deviation (SD) for continuous variables. We used a nonparametric rank-sum test for trend to examine any differences in the distribution of our study variables across the six months. All analyses were performed using Stata/MP version 15.1 software (StataCorp, 2015).

## RESULTS

### Frequency of COVID-19 infection among all VHA users

We identified a total of 43,267 VHA users with laboratory-confirmed COVID-19 during our study period. Of these cases, 5.4% (N=2,344) were diagnosed in March, 16.1% (N=6,947) in April, 10.6% (N=4,594) in May, 15.9% (N=6,867) in June, 33.2% (N=14,369) in July, and 18.8% (N=8,146) in August (P_trend_<0.001).

### Patient characteristics among VHA users hospitalized with COVID-19

Our cohort comprised 8,240 patients hospitalized with COVID-19 (Table 1). The average LOS was 10.2 days (SD=10.9), and 13.1% (N=1,081) experienced in-hospital mortality within 30-days of admission. The proportion of hospitalized patients increased overall (P_trend_<0.001) and fluctuated over time, with the highest observed in April (N=1,559, 18.9%) and July (N=2,279, 27.7%). The proportion of patients that experienced 30-day in-hospital mortality consistently declined and was three times lower in August (N=111/1,396, 8.0%) compared to March (N=221/890, 24.8%) (P_trend_<0.001).

**Table 1.** Patient characteristics* among 8,240 hospitalized Veterans Health Administration users with laboratory-confirmed coronavirus disease 2019 (COVID-19), March 1, 2020 – August 31, 2020

|  | March<br>N (%) | April<br>N (%) | May<br>N (%) | June<br>N (%) | July<br>N (%) | August<br>N (%) | Total<br>N (%) | P-trend |
| --- | --- | --- | --- | --- | --- | --- | --- | --- |
| **Total Hospitalized** | 890 (10.8) | 1,559 (18.9) | 889 (10.8) | 1,227 (14.9) | 2,279 (27.7) | 1,396 (16.9) | 8,240 (100.0) | <0.001 |
| Age in years |  |  |  |  |  |  |  | 0.001 |
| ≤49 | 78 (8.8) | 101 (6.5) | 68 (7.7) | 159 (13.0) | 255 (11.2) | 126 (9.0) | 787 (9.6) |  |
| 50-59 | 148 (16.6) | 171 (11.0) | 94 (10.6) | 181 (14.8) | 326 (14.3) | 179 (12.8) | 1,099 (13.3) |  |
| 60-69 | 235 (26.4) | 385 (24.7) | 250 (28.1) | 304 (24.8) | 591 (25.9) | 344 (24.6) | 2,109 (25.6) |  |
| 70-79 | 288 (32.4) | 515 (33.0) | 273 (30.7) | 390 (31.8) | 743 (32.6) | 475 (34.0) | 2,684 (32.6) |  |
| ≥80 | 141 (15.8) | 387 (24.8) | 204 (23.0) | 193 (15.7) | 364 (16.0) | 272 (19.5) | 1,561 (18.9) |  |
| Mean (SD) | 67.4 (13.0) | 70.7 (13.1) | 69.9 (13.5) | 66.4 (14.7) | 67.0 (13.8) | 68.4 (13.5) | 68.2 (13.7) | 0.001 |
| Male sex | 853 (95.8) | 1,490 (95.6) | 846 (95.2) | 1,148 (93.6) | 2,147 (94.2) | 1,315 (94.2) | 7,799 (94.7) | 0.01 |
| Race/Ethnicity |  |  |  |  |  |  |  | <0.001 |
| White | 313 (35.2) | 751 (48.2) | 465 (52.3) | 669 (54.5) | 1,242 (54.5) | 828 (59.3) | 4,268 (51.8) |  |
| Black | 508 (57.1) | 703 (45.1) | 354 (39.8) | 438 (35.7) | 813 (35.7) | 431 (30.9) | 3,247 (39.4) |  |
| Other | 69 (7.8) | 105 (6.7) | 70 (7.9) | 120 (9.8) | 224 (9.8) | 137 (9.8) | 725 (8.8) |  |
| Urban residence | 837 (94.0) | 1,426 (91.5) | 750 (84.4) | 1,018 (83.0) | 1,830 (80.3) | 954 (68.3) | 6,815 (82.7) | <0.001 |
| Priority Rating (1-8) |  |  |  |  |  |  |  | 0.56 |
| 1-4 | 266 (29.9) | 450 (28.9) | 260 (29.3) | 339 (27.6) | 649 (28.5) | 408 (29.2) | 2,372 (28.8) |  |
| 5-6 | 320 (36.0) | 577 (37.0) | 345 (38.8) | 509 (41.5) | 912 (40.0) | 526 (37.7) | 3,189 (38.7) |  |
| 7-8 | 304 (34.2) | 532 (34.1) | 284 (32.0) | 379 (30.9) | 718 (31.5) | 462 (33.1) | 2,679 (32.5) |  |
| Length of stay in days, Mean (SD) | 10.1 (10.9) | 10.8 (11.9) | 11.8 (12.4) | 10.6 (11.7) | 9.6 (9.9) | 9.3 (9.1) | 10.2 (10.9) | <0.001 |
| 30-day in-hospital mortality | 221 (24.8) | 325 (20.8) | 103 (11.6) | 115 (9.4) | 206 (9.0) | 111 (8.0) | 1,081 (13.1) | <0.001 |

**Note.** N=Number; SD=Standard Deviation.
\*Socio-demographic characteristics were examined at the time of hospitalization for COVID-19

Overall, hospitalized patients were older (M=68.2, SD=13,7), predominantly male (N=7,799, 94.7%), and white (N=4,268, 51.8%). The majority resided in an urban area (N=6,815, 82.7%), and close to 40% (N=3,189) had a priority rating of 5-6. The average age of hospitalized patients increased overall from March (M=67.4, SD=13.0) to August (M=68.4, SD=13.5), and patients aged 60 years and older consistently represented at least 70% of hospitalizations (P_trend_=0.001). Months with a greater proportion of younger (age <60 years) patients included March, June, and July. Black patients represented 57.1% (N=508/890) of hospitalizations in March, but consistently decreased to 30.9% (N=431/1,396) by August (P_trend_<0.001). There was a consistent decline in the proportion of hospitalized patients with urban residence from March (N=2,134/2,344, 91.0%) to August (N=5,529/8,146, 67.9%) (P_trend_<0.001).

### COVID-19 symptoms

The most frequently reported symptoms among hospitalized patients were fever (N=3,967, 48.1%), dyspnea (N=2,398, 29.1%), and cough (N=1,705, 20.7%) (Table 2). The proportion of patients reporting these symptoms significantly declined over time (P_trend_<0.001). Additionally, the proportion of patients without any record of symptoms more than tripled from March (N=86/890, 9.7%) to August (N=460/1,396, 33.0%) (P_trend_<0.001).

**Table 2.**
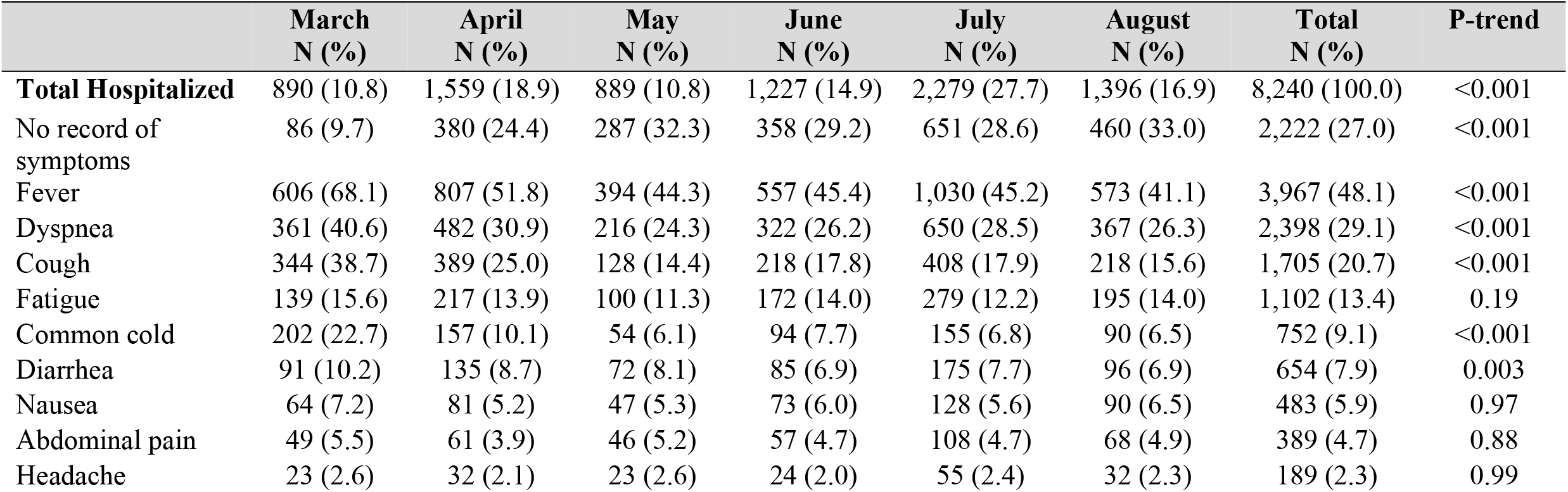
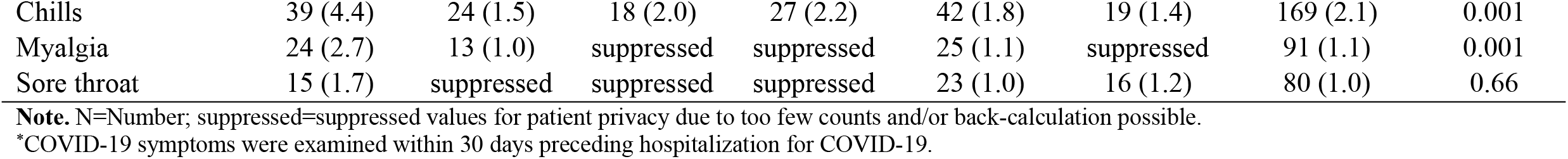
Patient symptoms* among 8,240 hospitalized Veterans Health Administration users with laboratory-confirmed coronavirus disease 2019 (COVID-19), March 1, 2020 – August 31, 2020

|  | March<br>N (%) | April<br>N (%) | May<br>N (%) | June<br>N (%) | July<br>N (%) | August<br>N (%) | Total<br>N (%) | P-trend |
| --- | --- | --- | --- | --- | --- | --- | --- | --- |
| **Total Hospitalized** | 890 (10.8) | 1,559 (18.9) | 889 (10.8) | 1,227 (14.9) | 2,279 (27.7) | 1,396 (16.9) | 8,240 (100.0) | <0.001 |
| No record of symptoms | 86 (9.7) | 380 (24.4) | 287 (32.3) | 358 (29.2) | 651 (28.6) | 460 (33.0) | 2,222 (27.0) | <0.001 |
| Fever | 606 (68.1) | 807 (51.8) | 394 (44.3) | 557 (45.4) | 1,030 (45.2) | 573 (41.1) | 3,967 (48.1) | <0.001 |
| Dyspnea | 361 (40.6) | 482 (30.9) | 216 (24.3) | 322 (26.2) | 650 (28.5) | 367 (26.3) | 2,398 (29.1) | <0.001 |
| Cough | 344 (38.7) | 389 (25.0) | 128 (14.4) | 218 (17.8) | 408 (17.9) | 218 (15.6) | 1,705 (20.7) | <0.001 |
| Fatigue | 139 (15.6) | 217 (13.9) | 100 (11.3) | 172 (14.0) | 279 (12.2) | 195 (14.0) | 1,102 (13.4) | 0.19 |
| Common cold | 202 (22.7) | 157 (10.1) | 54 (6.1) | 94 (7.7) | 155 (6.8) | 90 (6.5) | 752 (9.1) | <0.001 |
| Diarrhea | 91 (10.2) | 135 (8.7) | 72 (8.1) | 85 (6.9) | 175 (7.7) | 96 (6.9) | 654 (7.9) | 0.003 |
| Nausea | 64 (7.2) | 81 (5.2) | 47 (5.3) | 73 (6.0) | 128 (5.6) | 90 (6.5) | 483 (5.9) | 0.97 |
| Abdominal pain | 49 (5.5) | 61 (3.9) | 46 (5.2) | 57 (4.7) | 108 (4.7) | 68 (4.9) | 389 (4.7) | 0.88 |
| Headache | 23 (2.6) | 32 (2.1) | 23 (2.6) | 24 (2.0) | 55 (2.4) | 32 (2.3) | 189 (2.3) | 0.99 |
| Chills | 39 (4.4) | 24 (1.5) | 18 (2.0) | 27 (2.2) | 42 (1.8) | 19 (1.4) | 169 (2.1) | 0.001 |
| Myalgia | 24 (2.7) | 13 (1.0) | suppressed | suppressed | 25 (1.1) | suppressed | 91 (1.1) | 0.001 |
| Sore throat | 15 (1.7) | suppressed | suppressed | suppressed | 23 (1.0) | 16 (1.2) | 80 (1.0) | 0.66 |

**Note.** N=Number; suppressed=suppressed values for patient privacy due to too few counts and/or back-calculation possible.
\*COVID-19 symptoms were examined within 30 days preceding hospitalization for COVID-19.

### Clinical comorbidities among hospitalized and in-hospital deceased patients

Hospitalized patients had an average CCI score of 1.8 (SD=2.1), while those that experienced 30-day in-hospital mortality had an average CCI score of 2.3 (SD=2.3) (Table 3). CCI score significantly declined over time for both hospitalized (P_trend_<0.001) and in-hospital deceased (P_trend_=0.02) patients.

**Table 3.**
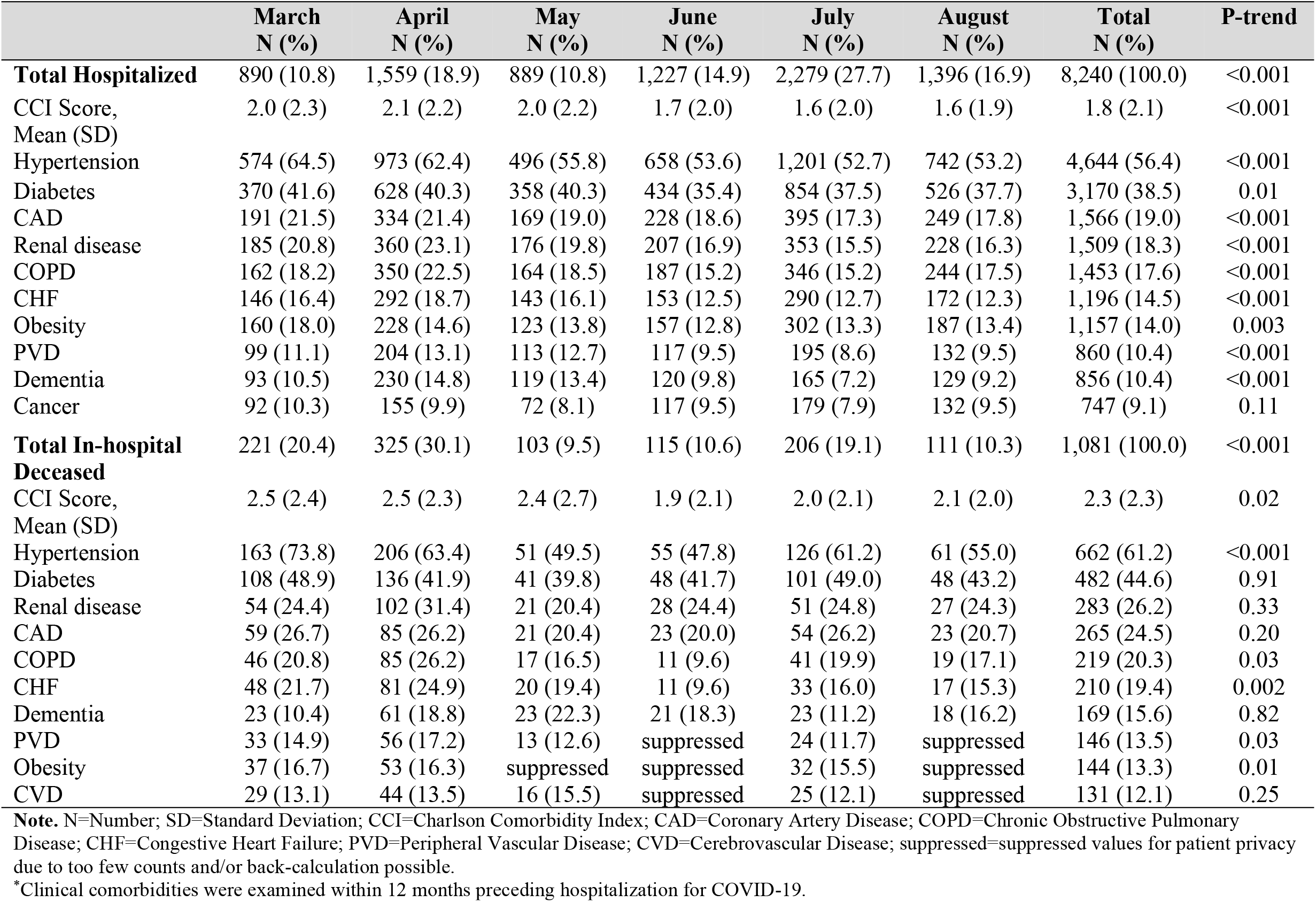
Top 10 clinical comorbidities* among hospitalized and in-hospital deceased Veterans Health Administration users with laboratory-confirmed coronavirus disease 2019 (COVID-19), March 1, 2020 – August 31, 2020

The top 10 comorbidities among hospitalized patients were hypertension (N=4,644, 56.4%), diabetes (N=3,170, 38.5%), coronary artery disease (N=1,566, 19.0%), renal disease (N=1,509, 18.3%), chronic obstructive pulmonary disease (N=1,453, 17.6%), congestive heart failure (N=1,196, 14.5%), obesity (N=1,157, 14.0%), peripheral vascular disease (N=860, 10.4%), dementia (N=856, 10.4%), and cancer (N=747, 9.1%). The proportion of hospitalized patients with these comorbidities significantly declined from March to August with the exception of cancer (P_trend_=0.11).

The top 10 comorbidities among in-hospital deceased patients were similar to those reported among hospitalized patients. These included hypertension (N=662, 61.2%), diabetes (N=482, 44.6%), renal disease (N=283, 26.2%), coronary artery disease (N=265, 24.5%), chronic obstructive pulmonary disease (N=219, 20.3%), congestive heart failure (N=210, 19.4%), dementia (N=169, 15.6%), peripheral vascular disease (N=146, 13.5%), obesity (N=144, 13.3%), and cerebrovascular disease (N=131, 12.1%).

Similarly, the proportion of in-hospital deceased patients with these comorbidities declined over time.

### Laboratory values among VHA users hospitalized with COVID-19

WBC, PC, and LC were low for 15.5% (N=1,280), 18.4% (N=1,517), and 41.9% (N=3,453) of hospitalized patients, respectively (Table 4). Over one-third of patients had elevated creatinine (N=3,153, 38.3%), BUN (N=2,804, 34.0%), AST (N=2,746, 33.3%), ferritin (N=3,614, 43.9%), and lactate (N=3,576, 43.4%). More than one-half (N=4,237, 51.4%) of patients had elevated CRP. For almost all laboratory values, patients hospitalized in March either had a similar or greater proportion of abnormal results compared to those hospitalized in later months. However, the proportion of patients with elevated CRP significantly increased over time (P_trend_<0.001).

**Table 4.** Laboratory values* among 8,240 hospitalized Veterans Health Administration users with coronavirus disease 2019 (COVID-19), March 1, 2020 – August 31, 2020

|  | March<br>N (%) | April<br>N (%) | May<br>N (%) | June<br>N (%) | July<br>N (%) | August<br>N (%) | Total<br>N (%) | P-trend |
| --- | --- | --- | --- | --- | --- | --- | --- | --- |
| **Total Hospitalized** | 890 (10.8) | 1,559 (18.9) | 889 (10.8) | 1,227 (14.9) | 2,279 (27.7) | 1,396 (16.9) | 8,240 (100.0) | <0.001 |
| WBC <4,000/μL | 151 (17.0) | 248 (15.9) | 119 (13.4) | 177 (14.4) | 363 (15.9) | 222 (15.9) | 1,280 (15.5) | 0.24 |
| Unknown/missing | 90 (10.5) | 131 (8.4) | 67 (7.5) | 94 (7.7) | 174 (7.6) | 122 (8.7) | 681 (8.3) |  |
| Absolute LC<br><1,000/μL | 438 (49.2) | 705 (45.2) | 362 (40.7) | 451 (36.8) | 916 (40.2) | 581 (41.6) | 3,453 (41.9) | 0.13 |
| Unknown/missing | 42 (4.7) | 81 (5.2) | 50 (5.6) | 95 (7.7) | 163 (7.2) | 104 (7.5) | 535 (6.5) |  |
| PC <150,000/μL | 194 (21.8) | 296 (19.0) | 132 (14.9) | 216 (17.6) | 429 (18.8) | 250 (17.9) | 1,517 (18.4) | 0.28 |
| Unknown/missing | 34 (3.8) | 72 (4.6) | 45 (5.1) | 74 (6.0) | 143 (6.3) | 91 (6.5) | 459 (5.6) |  |
| Creatinine >1.5<br>mg/dL | 380 (42.7) | 633 (40.6) | 333 (37.5) | 440 (35.9) | 874 (38.4) | 493 (35.3) | 3,153 (38.3) | <0.001 |
| Unknown/missing | 35 (3.9) | 68 (4.4) | 45 (5.1) | 71 (5.8) | 140 (6.1) | 90 (6.5) | 449 (5.5) |  |
| BUN >20 mg/dL | 335 (37.6) | 624 (40.0) | 343 (38.6) | 372 (30.3) | 691 (30.3) | 439 (31.5) | 2,804 (34.0) | <0.001 |
| Unknown/missing | 35 (3.9) | 68 (4.4) | 45 (5.1) | 71 (5.8) | 140 (6.1) | 90 (6.5) | 449 (5.5) |  |
| Total bilirubin ≥1.2<br>mg/dL | 47 (5.3) | 104 (6.7) | 41 (4.6) | 67 (5.5) | 164 (7.2) | 81 (5.8) | 504 (6.1) | 0.05 |
| Unknown/missing | 41 (4.6) | 82 (5.3) | 61 (6.9) | 90 (7.3) | 161 (7.1) | 111 (8.0) | 546 (6.6) |  |
| AST >40 U/L | 383 (43.0) | 606 (38.9) | 269 (30.3) | 342 (27.9) | 742 (32.6) | 404 (28.9) | 2,746 (33.3) | <0.001 |
| Unknown/missing | 49 (5.5) | 97 (6.2) | 68 (7.7) | 104 (8.5) | 181 (7.9) | 131 (9.4) | 630 (7.7) |  |
| ALT >40 U/L | 172 (19.3) | 226 (14.5) | 98 (11.0) | 175 (14.3) | 347 (15.2) | 200 (14.3) | 1,218 (14.8) | <0.001 |
| Unknown/missing | 47 (5.3) | 95 (6.1) | 63 (7.1) | 103 (8.4) | 180 (7.9) | 134 (9.6) | 622 (7.6) |  |
| Ferritin >300 ng/mL | 448 (50.3) | 754 (48.4) | 348 (39.2) | 491 (40.0) | 1,009 (44.3) | 564 (40.4) | 3,614 (43.9) | <0.001 |
| Unknown/missing | 265 (29.8) | 375 (24.1) | 230 (25.9) | 343 (28.0) | 600 (26.3) | 425 (30.4) | 2,238 (27.2) |  |
| Lactate >2.2<br>mmol/L | 465 (52.3) | 747 (47.9) | 342 (38.5) | 488 (39.8) | 987 (43.3) | 547 (39.2) | 3,576 (43.4) | <0.001 |
| Unknown/missing | 282 (31.7) | 455 (29.2) | 300 (33.8) | 416 (33.9) | 748 (32.8) | 540 (38.7) | 2,741 (33.3) |  |
| Troponin I ≥0.06<br>ng/mL | 148 (16.6) | 233 (15.0) | 105 (11.8) | 126 (10.3) | 297 (13.0) | 179 (12.8) | 1,088 (13.2) | 0.31 |
| Unknown/missing | 268 (30.1) | 482 (30.9) | 302 (34.0) | 349 (28.4) | 641 (28.1) | 457 (32.7) | 2,499 (30.3) |  |
| BNP >100 pg/mL | 216 (24.3) | 403 (25.9) | 222 (25.0) | 257 (21.0) | 471 (20.7) | 300 (21.5) | 1,869 (22.7) | 0.002 |
| Unknown/missing | 378 (42.5) | 734 (47.1) | 444 (49.9) | 572 (46.6) | 1,026 (45.0) | 714 (51.2) | 3,868 (46.9) |  |
| Procalcitonin >0.25 ng/mL | 212 (23.8) | 437 (28.0) | 188 (21.2) | 184 (15.0) | 373 (16.4) | 218 (15.6) | 1,612 (19.6) | <0.001 |
| Unknown/missing | 392 (44.0) | 670 (43.0) | 439 (49.4) | 596 (48.6) | 1,050 (46.1) | 713 (51.1) | 3,860 (46.8) |  |
| CRP >8.2 ng/mL | 441 (49.6) | 785 (50.4) | 441 (49.6) | 579 (47.2) | 1,233 (54.1) | 758 (54.3) | 4,237 (51.4) | <0.001 |
| Unknown/missing | 423 (47.5) | 696 (44.6) | 362 (40.7) | 545 (44.4) | 851 (37.3) | 508 (36.4) | 3,385 (41.1) |  |

**Note.** N=Number; WBC=White Blood Cell Count; LC=Lymphocyte Count; PC=Platelet Count; BUN=Blood Urea Nitrogen; AST=Aspartate Aminotransferase; ALT=Alanine Aminotransferase; BNP=Brain-type Natriuretic Peptide; CRP=C-reactive Protein.
\*Laboratory values were examined within seven days following hospitalization for COVID-19.

### Pharmacological and non-pharmacological COVID-19 treatment

Close to one-half (N=3,774, 45.8%) of hospitalized patients were treated with corticosteroids, the use of which significantly increased (P_trend_<0.001) (Table 5).

**Table 5.** Pharmacological and non-pharmacological treatment* among 8,240 hospitalized Veterans Health Administration users with coronavirus disease 2019 (COVID-19), March 1, 2020 – August 31, 2020

|  | March<br>N (%) | April<br>N (%) | May<br>N (%) | June<br>N (%) | July<br>N (%) | August<br>N (%) | Total<br>N (%) | P-trend |
| --- | --- | --- | --- | --- | --- | --- | --- | --- |
| **Total Hospitalized** | 890 (10.8) | 1,559 (18.9) | 889 (10.8) | 1,227 (14.9) | 2,279 (27.7) | 1,396 (16.9) | 8,240 (100.0) | <0.001 |
| **Pharmacological Treatments** |  |  |  |  |  |  |  |  |
| Anticoagulants | 799 (89.8) | 1,372 (88.0) | 792 (89.1) | 1,078 (87.9) | 2,098 (92.1) | 1,303 (93.3) | 7,442 (90.3) | <0.001 |
| Statins | 527 (59.2) | 894 (57.3) | 515 (57.9) | 693 (56.5) | 1,342 (58.9) | 796 (57.0) | 4,767 (57.9) | 0.78 |
| Antibiotics | 678 (76.2) | 931 (59.7) | 448 (50.4) | 588 (47.9) | 1,131 (49.6) | 692 (49.6) | 4,468 (54.2) | <0.001 |
| *Azithromycin* | 510 (75.2) | 514 (55.2) | 215 (48.0) | 280 (47.6) | 632 (55.9) | 349 (50.4) | 2,500 (56.0) | <0.001 |
| Antivirals | 529 (59.4) | 577 (37.0) | 232 (26.1) | 426 (34.7) | 946 (41.5) | 580 (41.6) | 3,290 (39.9) | 0.002 |
| *Hydroxychloroquine* | 492 (93.0) | 514 (89.1) | 27 (11.6) | suppressed | 19 (2.0) | suppressed | 1,074 (32.6) | <0.001 |
| *Remdesivir* | 10 (1.9) | 34 (5.9) | 191 (82.3) | 390 (91.5) | 874 (92.4) | 543 (93.6) | 2,042 (62.1) | <0.001 |
| Azithromycin +<br>Hydroxychloroquine | 370 (41.6) | 294 (18.9) | suppressed | suppressed | 14 (1.0) | suppressed | 698 (8.5) | <0.001 |
| NSAIDs | 455 (51.1) | 771 (49.5) | 470 (52.9) | 661 (53.9) | 1,239 (54.4) | 797 (57.1) | 4,393 (53.3) | <0.001 |
| Beta-blockers | 439 (49.3) | 738 (47.3) | 422 (47.5) | 530 (43.2) | 1,044 (45.8) | 700 (50.1) | 3,873 (47.0) | 0.83 |
| Corticosteroids | 245 (27.5) | 413 (26.5) | 214 (24.1) | 580 (47.3) | 1,440 (63.2) | 882 (63.2) | 3,774 (45.8) | <0.001 |
| *Dexamethasone* | 33 (13.5) | 43 (10.4) | 29 (13.6) | 391 (67.4) | 1,205 (83.7) | 732 (83.0) | 2,433 (64.5) | <0.001 |
| Bronchodilators | 397 (44.6) | 687 (44.1) | 348 (39.2) | 552 (45.0) | 1,043 (45.8) | 632 (45.3) | 3,659 (44.4) | 0.15 |
| ACE-inhibitors | 183 (20.6) | 315 (20.2) | 195 (21.9) | 296 (24.1) | 596 (26.2) | 343 (24.6) | 1,928 (23.4) | <0.001 |
| Vasopressors | 189 (21.2) | 256 (16.4) | 113 (12.7) | 123 (10.0) | 206 (9.0) | 119 (8.5) | 1,006 (12.2) | <0.001 |
| Immune-based<br>therapy | 55 (6.2) | 145 (9.3) | 45 (5.1) | 45 (3.7) | 93 (4.1) | 16 (1.2) | 399 (4.8) | <0.001 |
| *Tocilizumab* | 52 (94.5) | 133 (91.7) | 45 (100.0) | 45 (100.0) | 93 (100.0) | 16 (100.0) | 384 (96.2) | <0.001 |
| **Non-pharmacological Treatments** |  |  |  |  |  |  |  |  |
| Mechanical<br>ventilation | 279 (31.4) | 330 (21.2) | 150 (16.9) | 165 (13.5) | 354 (15.5) | 183 (13.1) | 1,461 (17.7) | <0.001 |
| Dialysis | 129 (14.5) | 165 (10.6) | 66 (7.4) | 69 (5.6) | 130 (5.7) | 59 (4.2) | 618 (7.5) | <0.001 |
| Supplemental oxygen | 67 (7.5) | 137 (8.8) | 57 (6.4) | 82 (6.7) | 178 (7.8) | 90 (6.5) | 611 (7.4) | 0.16 |

**Note.** N=Number; NSAIDs=Non-steroidal Anti-inflammatory Drugs; ACE=Angiotensin-converting Enzyme.
\*Receipt of any pharmacological and/or non-pharmacological treatment was examined during the course of hospitalization for COVID-19.

Dexamethasone represented 65% (N=2,433/3,774) of corticosteroids overall and increased from 14% (N=33/245) in March to 83% (N=732/882) by August (P_trend_<0.001). About two-fifths (N=3,290, 39.9%) of patients were treated with antivirals, which significantly declined overall (P_trend_<0.001). Among antivirals, Hydroxychloroquine was the most commonly used treatment in March (N=492/529, 93.0%) and April (N=514/577, 89.1%), while treatment with Remdesivir dramatically increased from May (N=191/232, 82.3%) to August (N=543/580, 93.6%). Very few patients were treated with Azithromycin in combination with Hydroxychloroquine (N=698, 8.5%). This was common in March (N=370/890, 41.6%), but dropped to less than 1% by August (P_trend_<0.001). Less than one-quarter (N=1,006, 12.2%) of patients were treated with vasopressors, which substantially declined over time (P_trend_<0.001).

Close to one-fifth (N=1,461, 17.7%) of hospitalized patients received mechanical ventilation, with greater representation in March (N=279/890, 31.4%) compared to August (N=183/1,396, 13.1%) (P_trend_<0.001). A small proportion of patients received dialysis (N=618, 7.5%) and supplemental oxygen (N=611, 7.4%) during their hospitalization for COVID-19. Treatment with dialysis declined from March (N=129/890, 14.5%) to August (N=59/1,396, 4.2%) (P_trend_<0.001), while use of supplemental oxygen remained constant over time (P_trend_=0.16).

## DISCUSSION

To the best of our knowledge, this is the first study that examines longitudinal trends in COVID-19 infection rates as well as clinical management within a national health care system. Within the VHA, COVID-19 cases and hospitalization rates fluctuated between March and August, but increased overall. We identified peaks for both cases and hospitalizations in April, July, and August. Patients hospitalized in March compared to later months were younger, mostly black, urban-dwelling, symptomatic, and more likely to present with abnormal laboratory findings. They also had worse overall health as indicated by higher CCI scores and frequency of baseline comorbidities. Lastly, we observed a decline in use of intensive (e.g., mechanical ventilation) and experimental (e.g., Hydroxychloroquine) treatments for COVID-19, as well as a consistent decline in 30-day in-hospital mortality over time.

Many prior studies have worked to identify risk factors for COVID-19 infection, severity, and fatality.^1,3–18^ Literature published early on in the pandemic, as well as more recent studies, have consistently reported an elevated risk of both testing positive and severe or fatal disease among older populations (age ≥65 years) after adjusting for sociodemographic characteristics and comorbidities.^4,9,13,16^ In our study, we found that the majority of patients hospitalized with COVID-19 were older. However, we also identified a higher proportion of younger patients hospitalized in months with corresponding elevations in infection rates. This finding may support the notion that young and presumably healthy individuals can contribute to the inadvertent spread of COVID-19.^9,17,19^ Therefore, containing disease spread may require more ubiquitous government restrictions and policies including testing, contact tracing, social distancing and work-from-home guidelines.^9,23^ Moreover, we identified a significant decline in 30-day in-hospital mortality and treatment-related indicators of severe illness (e.g., receipt of mechanical ventilation or vasopressors) from March to August despite an overall increase in the average age of hospitalized patients.^13^ This phenomenon may be due to experience gained from treating patients with COVID-19 and/or effective action taken by the VHA to meet care and resource demands, as evidenced by the positive transition in treatment modalities. During our study period, we observed treatment with experimental and ineffective drug regimens such as Hydroxychloroquine decline to less than 1% by August, while use of more evidence-backed treatments such as Dexamethasone and Remdesivir significantly increased.^11,19,33,34^

In addition to older age, prior work has documented evidence of increased risk for COVID-19 infection, severity, and/or fatality among patients with certain symptoms, pre-existing medical conditions, and abnormal laboratory findings.^4,7,13,14,28^ Although the significance and magnitude of associations varied between studies, our finding of declining mortality seems to support the idea of greater risk for severe or fatal disease among patients that have these risk factors. In our study, patients hospitalized in March compared to later months had more exacerbating symptoms (e.g., fever, dyspnea, cough) and evidence of biomarkers on inflammation, infection, cardiac and muscle injury, decreased liver and kidney function.^4,7,13,14,28^ Additionally, we found that both hospitalized and deceased patients had more underlying comorbidities in March compared to later months including severe cardiovascular diseases, diabetes, renal disease, and obesity. However, deceased patients had a higher frequency of such comorbidities. While more comorbidities may contribute to disease severity or fatality, they may not contribute to greater risk for COVID-19 infection as we did not find proportional declines in cases or hospitalizations. The specific reasons for the apparent decline in case severity and fatality are unclear, but may be due to increased dissemination of risk information and potentially greater compliance with public health measures among those with poorer overall health.

Lastly, various studies have reported elevated hospitalization and mortality rates among black compared to white patients with COVID-19, citing differences in access and quality of care, more underlying comorbidities, and lower SES as potential reasons for these disparities.^6,8,10,35–37^ In our study, we observed a higher proportion of black compared to white patients hospitalized with COVID-19 in March, similar proportions of black and white patients in April, and substantially fewer black patients from May through August. However, black patients consistently had a hospitalization rate that was more than two times their overall veteran population share of 12%.^38^ The initially higher proportion of black patients hospitalized with COVID-19 in our study could potentially be explained by structural confounding in which more black patients reside in densely populated urban areas that were also more likely to be affected during the first outbreak of COVID-19.^5,13,36^ Additionally, more black patients may work in service and essential industries that do not allow work from home and create challenges for social distancing.^5,39^ Although we did not explore differences in mortality by race, a recent study by Rentsch et al. on national cohort of 16,317 VHA patients with COVID-19 found evidence of greater adjusted risk for contracting COVID-19 among racial and ethnic minorities, but no difference in 30-day mortality.^5^ Unlike the general U.S. population, all patients treated within the VHA have insurance coverage, and could readily seek high quality health care and treatment services if needed.^5,13,40^ Therefore, in addition to bolstering the capacity of health care systems and implementation of strict public health measures, ensuring equitable care access should be a central goal of COVID-19 mitigation efforts.^5,9,13^

### Limitations

We acknowledge important limitations to our study. The VHA typically treats a population that is older, medically complex, and have greater risk behaviors compared to the general U.S. population.^41^ However, prior work has found no evidence of differences in comorbidity burden between VHA and non-VHA users after controlling for age, sex, race, geographic region, and urbanicity of residence.^24^ Additionally, the frequency and type of comorbidities identified in this study are similar to those reported among non-VHA patients hospitalized with COVID-19.^15–17^ Therefore, our findings may still be generalizable to the larger U.S. population. Our analysis was limited to patients diagnosed and hospitalized for COVID-19 within the VHA health care system and may not capture cases, hospitalizations, treatments received, or deaths occurring at non-VHA facilities. We reported the month-to-month and cumulative disease burden on the VHA, which may not account for regional differences in sociodemographic characteristics, government restrictions and policies, or the magnitude and timing of case peaks. Future work on the regional intensity and timing of outbreaks may be beneficial for identifying specific areas with elevated disease burden and help inform allocation of limited resources.

## CONCLUSIONS

Our study provides greater insight on the longitudinal trends in COVID-19 infection rates and clinical management within a national health care system. We found evidence of declining COVID-19 severity and 30-day in-hospital mortality, despite increases in cases, hospitalizations, and average age. Therefore, actions taken by the VHA may have been effective for bolstering treatment capacity and mitigating disease burden at the patient- and health system-level.

